# Whole genome assembly and annotation of the King Angelfish (*Holacanthus passer*) gives insight into the evolution of marine fishes of the Tropical Eastern Pacific

**DOI:** 10.1101/2023.11.08.566026

**Authors:** Remy Gatins, Carlos F. Arias, Carlos Sánchez, Giacomo Bernardi, Luis F. De León

**Author notes:** Corresponding author: Remy Gatins.

## Abstract

*Holacanthus* angelfishes are some of the most iconic marine fishes of the Tropical Eastern Pacific (TEP). However, very limited genomic resources currently exist for the genus. In this study we: i) assembled and annotated the nuclear genome of the King Angelfish (*Holacanthus passer),* and ii) examined the demographic history of *H. passer* in the TEP. We generated 43.8 Gb of ONT and 97.3 Gb Illumina reads representing 75X and 167X coverage, respectively. The final genome assembly size was 583 Mb with a contig N50 of 5.7 Mb, which captured 97.5% complete Actinoterygii Benchmarking Universal Single-Copy Orthologs (BUSCOs). Repetitive elements account for 5.09% of the genome, and 33,889 protein-coding genes were predicted, of which 22,984 have been functionally annotated. Our demographic model suggests that population expansions of *H. passer* occurred prior to the last glacial maximum (LGM) and were more likely shaped by events associated with the closure of the Isthmus of Panama. This result is surprising, given that most rapid population expansions in both freshwater and marine organisms have been reported to occur globally after the LGM. Overall, this annotated genome assembly will serve as a resource to improve our understanding of the evolution of *Holacanthus* angelfishes while facilitating novel research into local adaptation, speciation, and introgression in marine fishes.

## Introduction

The King angelfish, *Holacanthus passer*, is one of the most iconic fish species of the Tropical Eastern Pacific (TEP) (Figure 1). Its distribution ranges from the Northern Gulf of California (Sea of Cortez) to Peru, including the Revillagigedos, Cocos, Malpelo, and the Galápagos Islands [1,2] (Figure 1C). Due to their conspicuous coloration, the King angelfish have become a target for the aquarium trade [2], with individuals costing between $150 and $900 (at the time of publication), while individuals of the sister species, *H. clarionensis*, endemic to the Revillagigedos, have sold for up to $15,000. *Holacanthus passer* is currently protected under the conservation regulation in Mexico (Norma Official Mexicana) [2], but is identified as having a stable population under the IUCN red list [3]. *Holacanthus* angelfishes are protogynous sequential hermaphrodites, changing sex from female to male as they grow. They exhibit sexual dimorphism (pelvic fin coloration) (Figure 1A) [4], and can partition their habitat by sex and size classes [5]. They are important sponge feeders and herbivores but have also been observed feeding in the water column on fish feces [2,5] and interacting as fish cleaners [6]. Additionally, their social organization can vary from solitary individuals to harems [4].

**Figure 1.**
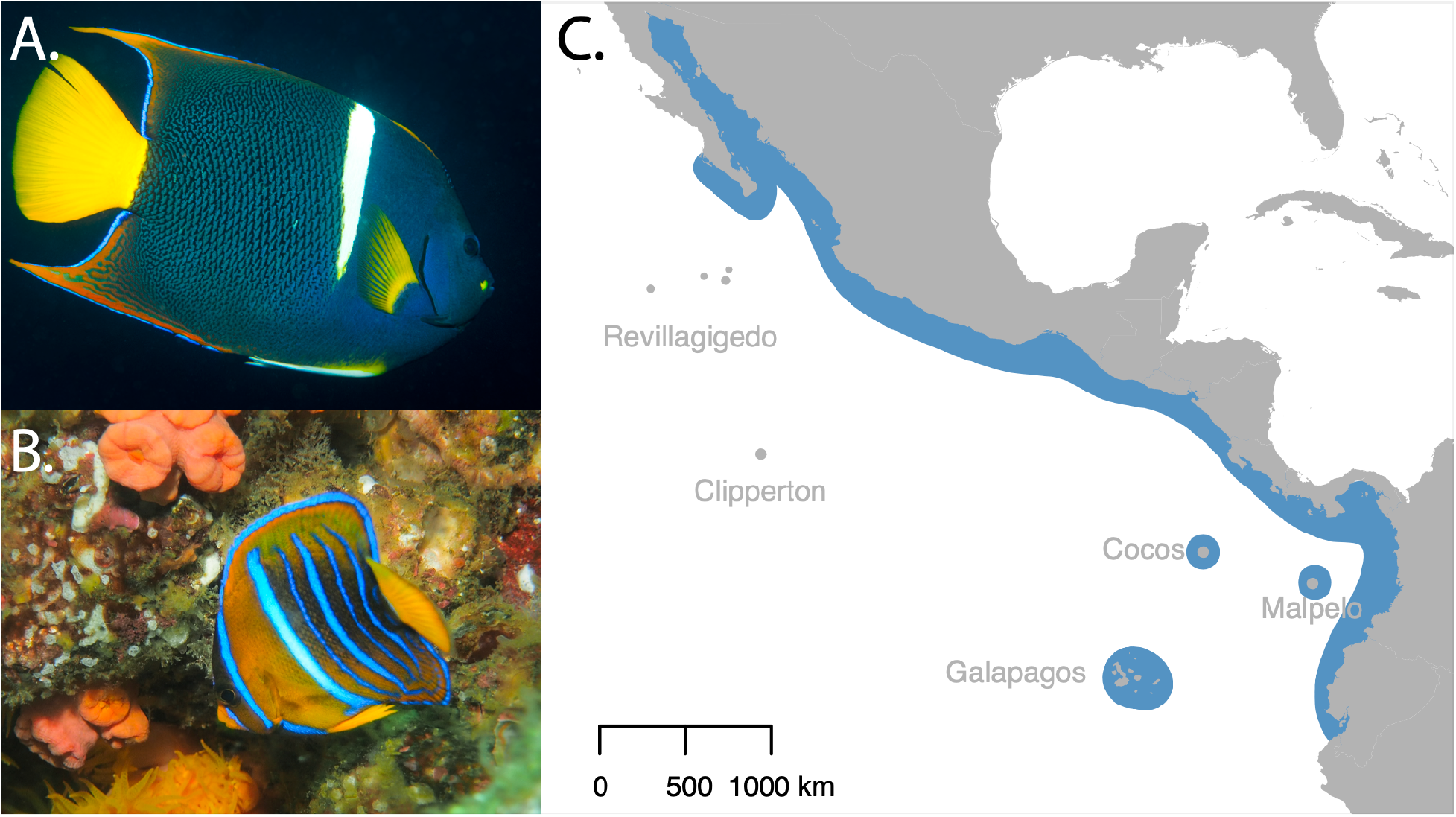
Pictures of an adult male (A) and juvenile (B) King angelfish, *Holacanthus passer.* Males can be identified by their white pelvic fin. Panel C shows a map *H. passer*’s range distribution in blue across the Tropical Eastern Pacific. Photo credits: Remy Gatins.

The genus *Holacanthus* is an interesting model system for assessing the drivers of diversification in marine fishes. Although it comprises of only seven species, it presents a complex history of diversification, which includes three modes of speciation: allopatric, peripatric, and sympatric [7,8]. Following the closure of the Isthmus of Panama around 3.2 to 2.8 Mya [9], two clades of *Holacanthus* were separated in the Atlantic and Pacific Oceans by the newly formed Isthmus. These so-called geminate species [10] are estimated to have diverged allopatrically approximately 1.7 to 1.4 Mya [7,8,11] along with about 40 other marine fishes (Jordan 1908; Thacker 2017) and many invertebrates [12,13]. Within each ocean basin, additional *Holacanthus* species diverged approximately 1.5 Mya. The Tropical Eastern Pacific (TEP) clade, which consists of *H.* passer, *H. limbaughi*, and *H. clarionensis,* is thought to have diverged via peripatry. In contrast, the Tropical Western Atlantic (TWA) clade, comprised by *H. bermudensis* and *H. ciliaris,* is thought to have diverged in sympatry [7,8]. The last two *Holacanthus* species, *H. tricolor* and *H. africanus*, are considered the sister taxon of the TEP-TWA clade, and the most ancestral *Holacanthus* taxon, respectively.

The increased accessibility of novel genomic tools has led to a rapid proliferation of whole-genome assemblies for non-model species. Recent genome assembly studies have used of a combination of short and accurate (∼99%) Illumina data with long, but less accurate, single-reads (∼95%) generated by Oxford Nanopore (ONT) or PacBio sequencing [14–18]. Hybrid assemblies can deliver real-time targeted sequencing, while improving genome assembly contiguity and completeness [14–16,19]. Thus, to facilitate the study of the history of diversification in *Holacanthus* and the interesting evolutionary dynamics associated with the closure of the Isthmus of Panama in the TEP, here we: i) deliver a high-quality whole genome assembly of the King Angelfish, *Holacanthus passer;* and ii) examined the demographic history of *H. passer* in the TEP using *de novo* genome sequence data.

## Main Content

### Context

#### Genome assembly

The final assembled and polished genome of *Holacanthus passer* yielded a total size of ∼583 Mb gathered in 486 contigs, with the largest contig at 17 Mb and a contig N50 of 5.7 Mb (Table 1). The 476 sequence fragments that make up the assembly contain zero gaps and are therefore described as contigs instead of scaffolds throughout the text. The final assembly was slightly larger than the initial ∼579 Mb estimated by GenomeScope (Figure 2A) as well as the initial 581 Mb assembly before the polishing iterations. Kraken identified approximately 100 kb of potential contaminants, none of which were identified using Blobtools (Figure 2B) and were thus retained in the assembly. Detailed assembly statistics after the first initial assembly and consecutive polishing rounds can be found in Table 1. The number of contigs remained at 486 contigs throughout the assembly. After four iterations of polishing using ONT and Illumina reads, BUSCO completeness improved from 82.4% to 97.5% for the Actinopterygii dataset (n = 4,584) and 90.1% to 95.4% in the Eukaryota dataset (n = 303). The largest completeness increase (10.6%) in the BUSCO Actinopterygii dataset occurred after the first ONT polishing iteration, while in the Eukaryota dataset the highest increase (2.3%) occurred after the first ONT polishing and the second Illumina polishing (Table 1). Additionally, the N50 contig length increased from 5.6 to 5.7 Mb after polishing. These results indicate that polishing with both ONT and Illumina reads greatly improved the assembly, by correcting assembly bases, fixing misassemblies, and filling assembly gaps. Moreover, contiguity did not improve after the initial assembly carried out with the Wtdbg2 assembler using long ONT reads. This suggests that the assembler and initial input reads play an important role in how contiguous the assembled genome will be, while multiple polishing iterations will further improve upon the accuracy of the assembly.

**Figure 2.**
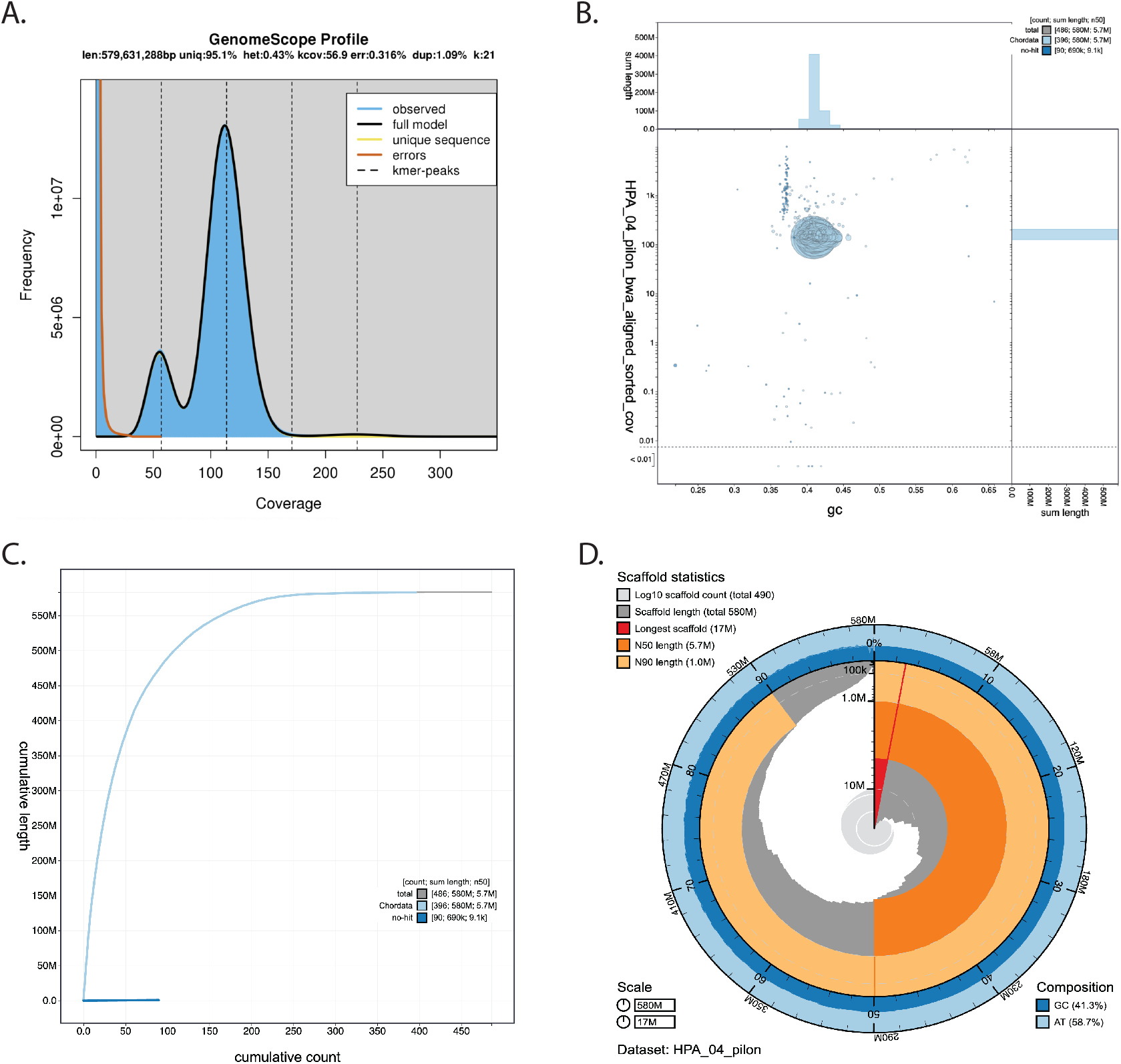
Assembly summary statistics views of of *Holacanthus passer* using GenomeScope VX (A) and Blobtoolkit Viewer (B-D). (A) Histogram for the 21 k-mer distribution of Illumina short reads. The highest frequency of k-mer coverage was seen around 110X (excluding k-mers with low coverage). (B) Blob plot showing the distribution of assembly scaffolds based on GC proportion and coverage. Circles are colored by phylum and circle size is relative to the number of sequence length. (C)Cumulative assembly plot showing curves of subsets of scaffolds assigned to each phylum relative to the overall assembly. (D) Snail plot summary of the assembly statistics.

**Table 1.** Step-by-step genome assembly and annotation statistics of the King angelfish (Holacanthus passer).

| Genome assembly | Nanopore |  |  | Nanopore + Illumina |  |
| --- | --- | --- | --- | --- | --- |
|  | Wtdbg2 | Wtdbg2 + 1X<br>Racon | Wtdbg2 + 2X<br>Racon | Wtdbg2 + 2X<br>Racon + 1X Pilon | Wtdbg2 + 2X<br>Racon + 2X Pilon |
| Total assembly size of contigs (bp) | 581 422 425 | 583 574 933 | 583 552 491 | 583 601 337 | 583 528 366 |
| Number of contigs | 486 | 486 | 486 | 486 | 486 |
| N50 contig length (bp) | 5 681 869 | 5 707 473 | 5 709 778 | 5 708 674 | 5 708 022 |
| N90 contig length (bp) | 997 074 | 1 000 168 | 1 000 597 | 1 000 715 | 1 000 532 |
| Longest contig (bp) | 17 088 287 | 17 147 963 | 17 147 963 | 17 150 647 | 17 148 928 |
| GC % |  |  |  |  | 41.27 |
| **Actinopterygii** |  |  |  |  |  |
| Complete BUSCOs | 3779 (82.4%) | 4263 (93%) | 4296 (93.7%) | 4468 (97.5%) | 4471 (97.5%) |
| Complete and single-copy BUSCOs | 3674 (80.1%) | 4133 (90.2%) | 4163 (90.8%) | 4364 (95.2%) | 4368 (95.3%) |
| Complete and duplicated BUSCOs | 105 (2.3%) | 130 (2.8%) | 133 (2.90%) | 104 (2.3%) | 103 (2.2%) |
| Fragmented BUSCOs | 374 (8.2%) | 176 (3.8%) | 155 (3.40%) | 38 (0.8%) | 37 (0.8%) |
| Missing BUSCOs | 431 (9.4%) | 145 (3.2%) | 133 (2.9%) | 78 (1.7%) | 76 (1.7%) |
| **Eukaryota** |  |  |  |  |  |
| Complete BUSCOs | 273 (90.1%) | 280 (92.4%) | 280 (92.4%) | 282 (93.10%) | 289 (95.4%) |
| Complete and single-copy BUSCOs | 267 (88.1%) | 270 (89.10%) | 270 (89.10%) | 267 (88.1%) | 274 (90.4%) |
| Complete and duplicated BUSCOs | 6 (2.0%) | 10 (3.3%) | 10 (3.3%) | 15 (5%) | 15 (5%) |
| Fragmented BUSCOs | 4 (1.3%) | 3 (1%) | 4 (1.3%) | 2 (0.7%) | 2 (0.7%) |
| Missing BUSCOs | 26 (8.6%) | 20 (6.60%) | 19 (6.3%) | 19 (6.2%) | 12 (3.9%) |
| **Annotation** |  |  |  |  |  |
| Number of protein-coding genes |  |  |  |  | 33 793 |
| Mean gene length (bp) |  |  |  |  | 10 807 |
| Number of CDSs |  |  |  |  | 388 693 |
| Longest gene (bp) |  |  |  |  | 315 502 |
| Functionally annotated |  |  |  |  | 22 992 |

The King angelfish genome assembly presented here is comparable in quality to other recently published fish genomes. Available genome assemblies of the most closely related fish species, such as, *Centropyge vrolikii* [18] and *Chaetodon austriacus* [20], exhibit slightly larger genome sizes of 696.5 Mb and 712.2 Mb, respectively. Importantly, our King angelfish genome resulted in a much more contiguous assembly (*H. passer*: 450; *C. vrolikii*: 30,500; *C. austriacus*: *13*,441) and a N50 of 5.7 Mb that is smaller than the N50 of *C. vrolikii* (9 Mb), but larger than that of *C. austriacus* (0.17 Mb) (Table 2). Regarding genome completeness, *H. passer* showed a slightly higher number of complete orthologous matches in BUSCO using the Actinopterygii (odb9) dataset than *C. vrolikii* and *C. austriacus* (Figure 3). When compared with numerous other recently published chromosome level fish genomes, *H. passer* showed comparable, if not higher, BUSCO scores despite not being a chromosome level assembly (Figure 3). In general, our assembly is highly contiguous with zero gaps, which could result in less fragmented genes. Overall, this *H. passer* assembly will serve as a high-quality genomic reference assembly for the Pomacanthidae family, and it exemplified how N50 values do not always correlate with the best BUSCO scores as outlined in Jauhal and Newcomb [21].

**Figure 3.**
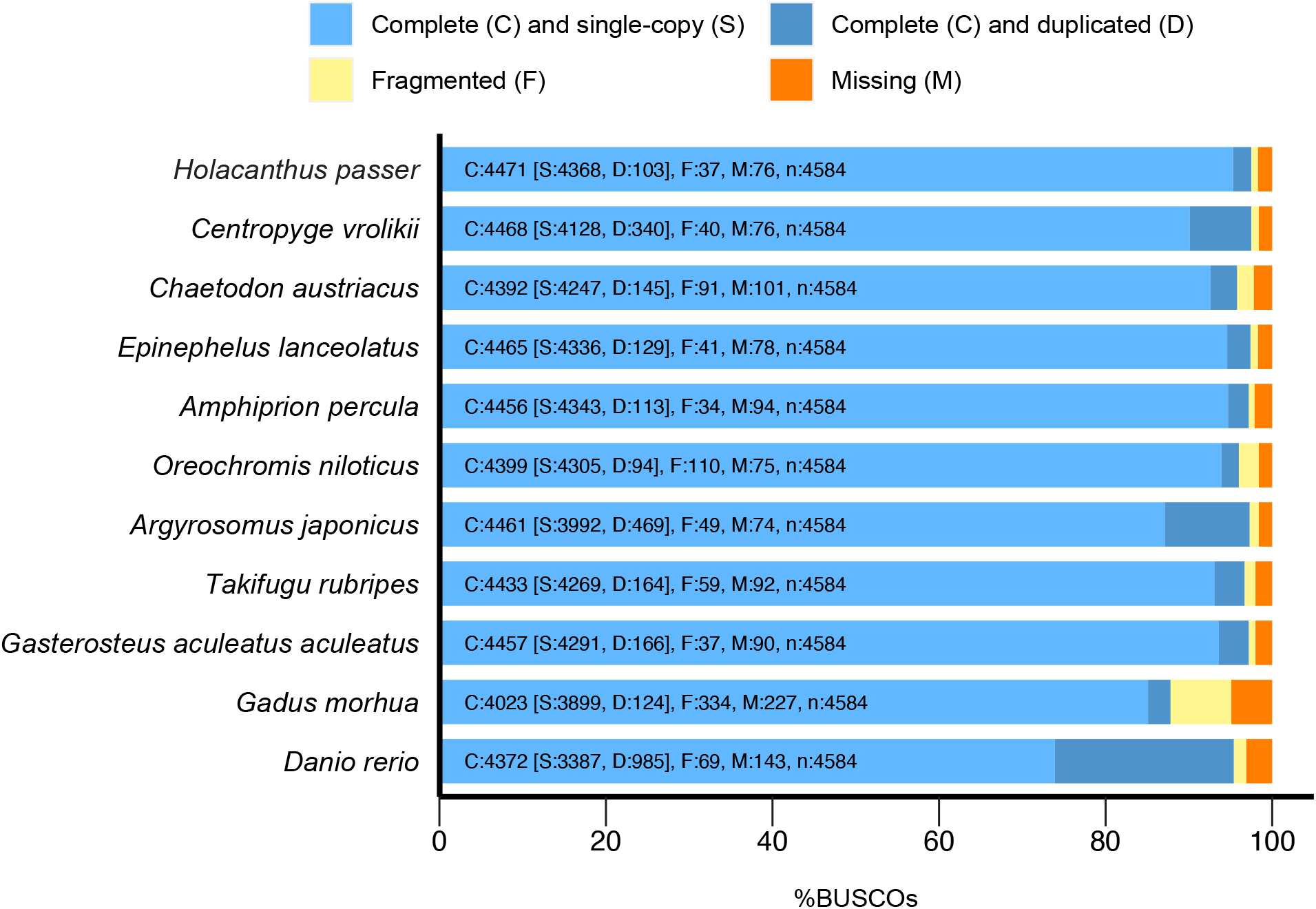
Comparative analysis of genome assembly completeness. BUSCO completeness of the *Holacanthus passer* genome assembly (first row) assessed by the 4584 orthologous actynopterygii (odb9) dataset. For comparison, we also assessed BUSCO scores for two closely related species (Pomacanthidae family: *C. vrolikii*, *C. austriacus*) and eight not closely related fish genomes to compare assemblies across fish biodiversity.

**Table 2.**
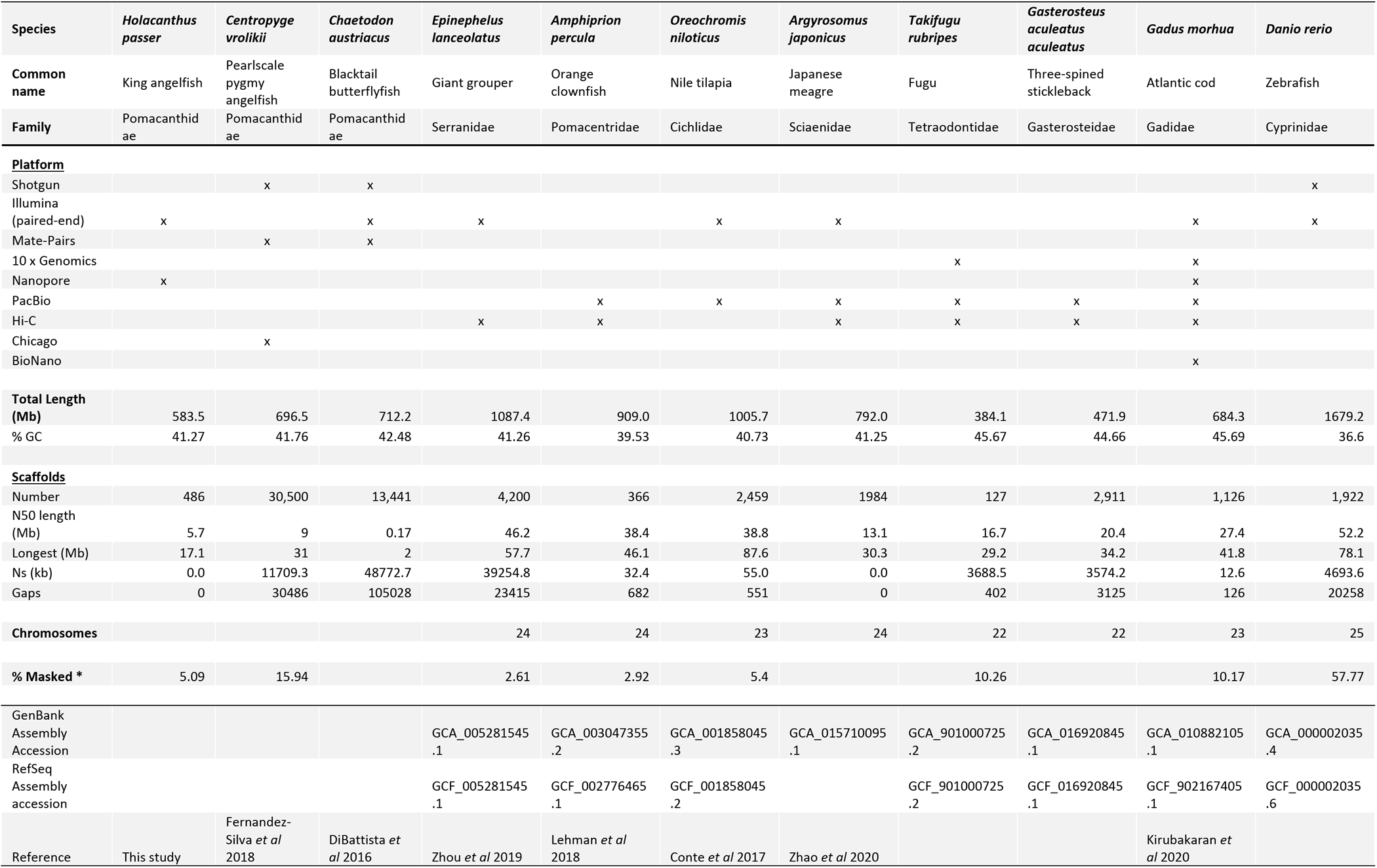
Comparison summary statistics for 11 selected fish genome assemblies, including *Holacanthus passer* from this study.

#### Genome annotation

RepeatMasker estimated that 5.09% of the genome consisted of repetitive sequences, primarily LINEs (0.85%), LTR elements (0.31%), DNA transposons (1.36%) and simple repeats (2.14%) (Table 3). Repeat content was nearly identical to that estimated by GenomeScope (4.9%). GeMoMa identified 33 793 gene models and 388 693 CDSs, where 67.8% (22 992) of the gene models had a functional annotation (Table 1). The number of coding sequences identified for *H. passer* was within the range of those found in other closely related fish species genomes (see https://www.ncbi.nlm.nih.gov/genome/annotation_euk/all/; assembled and annotated fish genomes, visited April 28, 2021).

**Table 3.** Summary output of repetitive elements of *H. passer* predicted by RepeatMasker v. 2.9.0+. The query species was assumed to be *Danio rerio*.

|  |  |
| --- | --- |
| **sequences:** | 486 |
| **total length:** | 583528366 bp (583528366 bp excl N/X-runs) |
| **GC level:** | 41.27% |
| **bases masked:** | 29714081 bp (5.09 %) |

|  | Number of<br>elements* | Length<br>occupied (bp) | Percentage of<br>sequence |
| --- | --- | --- | --- |
| **Retroelements** | 32172 | 6939127 | 1.19% |
| SINEs: | 1265 | 127915 | 0.02% |
| Penelope | 303 | 35535 | 0.01% |
| LINEs: | 19022 | 4977480 | 0.85% |
| CRE/SLACS | 0 | 0 | 0.00% |
| L2/CR1/Rex | 13025 | 3278888 | 0.56% |
| R1/LOA/Jockey | 644 | 120329 | 0.02% |
| R2/R4/NeSL | 299 | 122053 | 0.02% |
| RTE/Bov-B | 1556 | 536242 | 0.09% |
| L1/CIN4 | 2571 | 723359 | 0.12% |
| LTR elements: | 11885 | 1833732 | 0.31% |
| BEL/Pao | 1085 | 311809 | 0.05% |
| Ty1/Copia | 25 | 16958 | 0.00% |
| Gypsy/DIRS1 | 6190 | 1075740 | 0.18% |
| Retroviral | 2370 | 223106 | 0.04% |
| **DNA transposons** | 67101 | 7958272 | 1.36% |
| hobo-Activator | 24022 | 2283863 | 0.39% |
| Tc1-IS630-Pogo | 10113 | 2729978 | 0.47% |
| En-Spm | 0 | 0 | 0.00% |
| MuDR-IS905 | 0 | 0 | 0.00% |
| PiggyBac | 217 | 31373 | 0.01% |
| Tourist/Harbinger | 2025 | 231198 | 0.04% |
| Other (Mirage,<br>P-element, Transib) | 1538 | 262794 | 0.05% |
| Rolling-circles | 421 | 48093 | 0.01% |
| Unclassified: | 269 | 71601 | 0.01% |
| Total interspersed<br>repeats: |  | 14969000 | 2.57% |
| Small RNA: | 1676 | 161165 | 0.03% |
| Satellites: | 961 | 80911 | 0.01% |
| Simple repeats: | 303686 | 12479070 | 2.14% |
| Low complexity: | 38530 | 2144093 | 0.37% |

#### Demographic history of H. passer

The demographic history analysis of *H. passer* showed two extreme scenarios (Figure 4). When considering a faster mutation rate (μ) of 10^−8^, the population showed a slow expansion ∼300 Kya, with a small population decline occurring ∼70 Kya, followed by a second rapid expansion 30 Kya that reached a maximum effective population size of ∼300,000 individuals (Figure 4A). When using a slower mutation rate of 10^−9^, the population showed an initial expansion around 2.8 Mya, with a small decline ∼600 Kya, and the subsequent rapid expansion 300 Kya, reaching a maximum effective population size of ∼2,800,000 individuals (Figure 4B).

**Figure 4.**
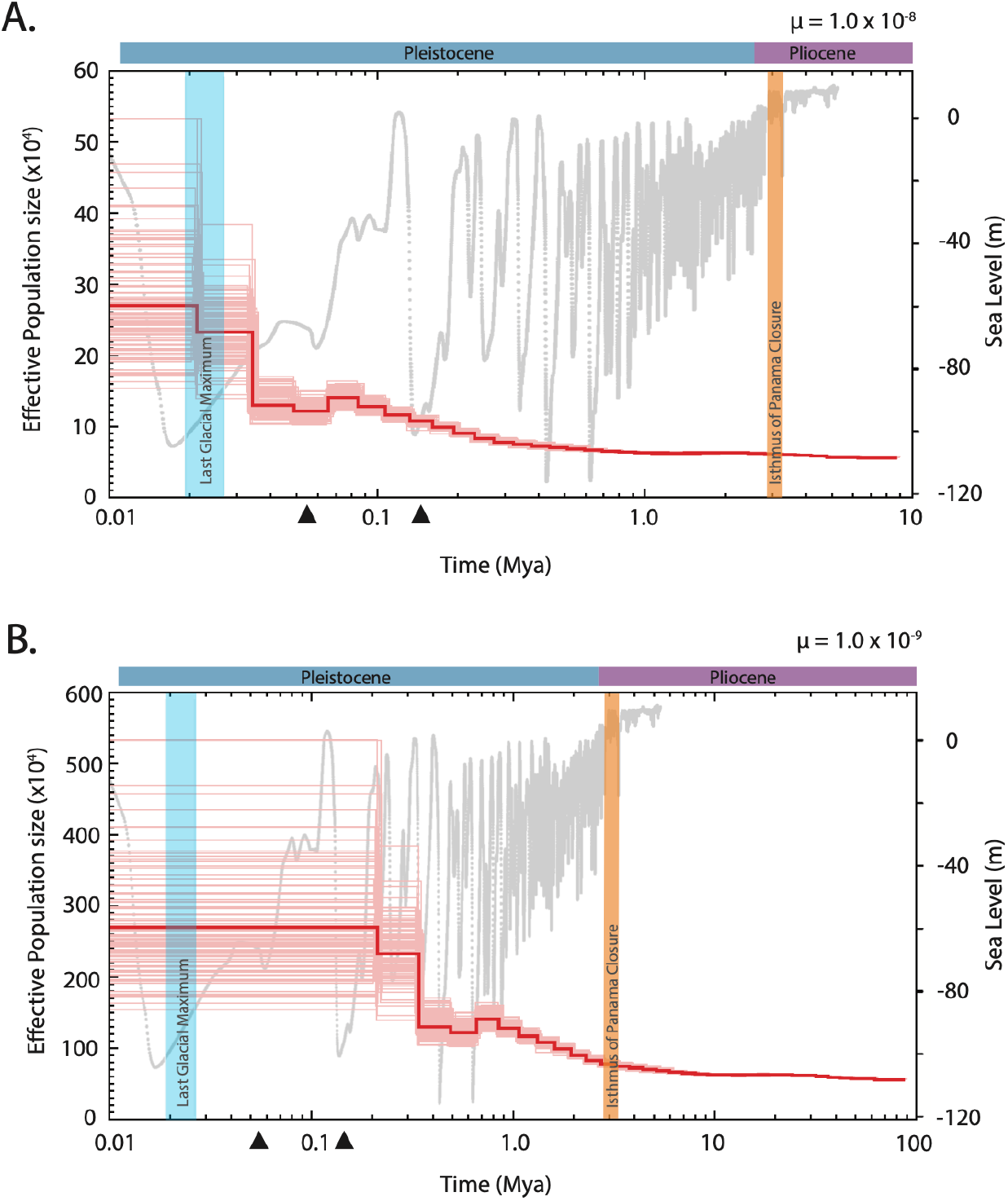
Genome-wide demographic history in *Holacanthus passer*. PSMC analysis showing the demographic history (red line) of *H. passer* using a generation time of 5 years and a mutation rate (μ) of 10-8 (A) and 10-9 (B). Global sea level model fluctuations over the past 5 million years are shown in the background (grey) [data from 50]. Vertical blue bars refer to the last glacial maximum (LGM) period (∼19-26.5 kya) and the orange bar represents the closure of the Isthmus of Panama (∼3.2-2.8 Mya). Triangles represent marine population expansion events previously recorded in the Tropical Eastern Pacific (see text).

Considering the slower mutation rate scenario, an effective population size in the order of millions of individuals for *H. passer* seems plausible when you consider the vast available habitat it occupies compared to its sister species *H. limbaughi* who’s effective population size was estimated ∼60,000 individuals [22]. *H. limbaughi* is endemic to Clipperton Island and occupies a fraction of the distribution of *H. passer*, which is found across the entire TEP coastline. However, considering the higher mutation rate scenario may seem likely when observing the first rapid population expansion occurring much after the closure of the Isthmus of Panama once oceanographic conditions in the TEP became more suitable.

*H. passer* was previously estimated to have diverged from its geminate Atlantic species (*H. ciliaris*) between 1.7 and 1.4 Mya [7,8], considering a molecular clock that was calibrated according to the closure of the Isthmus of Panama dated around 3.1 to 3.5 Mya [11]. However, recent studies suggest the closure of the Isthmus of Panama might have happened more recently, around 2.8 Mya [9]. Therefore, the genetic divergence between Holacanthus geminates could be more recent than previously believed.

After the closure of the Isthmus, oceanographic conditions in the TEP varied drastically following sea level changes due to multiple glaciation periods in the Pleistocene [23,24], likely leading to important demographic consequences [25]. Most rapid population expansions in both freshwater [26] and marine organisms [27] have been reported to occur globally after the last glacial maximum (LGM) that took place from 26.5 to 19 Kya [28]. However, only a few species have reported population expansions prior to the LGM [27]. On the contrary, in the TEP, most studies that have assessed the demographic history of marine organisms have found population expansions that precede the LGM [29–32] and few reporting population expansions in the last 20 Kya [32,33]. For instance, the goby, *Elacatinus puncticulatus,* and clingfish, *Gobieosox adustus,* experienced a population expansion around 170-130 Kya and 200-150 Kya, respectively [29,31]. While another reef fish, *Anisotremus interruptus*, experienced an expansion in its continental populations after the LGM (∼5 kya). Interestingly, *A. interruptus* populations from the oceanic islands of Revillagigedos and the Galapagos Archipelago showed earlier expansions at around 55 kya [32]. Yet, all demographic history studies in the TEP to date are based on single mitochondrial markers.

To the best of our knowledge, our study is the first to assess the demographic history of a marine fish in the TEP using genome-wide nuclear DNA. Our results support previous findings of marine population expansions in the TEP occurring prior to the LGM [29–32]. This pattern is consistent with our analyses using both slow and fast mutation rates for *H. passer*, which showed population expansions beyond 30 Kya. Overall, drops in sea level decrease the available marine habitat, potentially restricting gene flow between populations, thus resulting in population bottlenecks. This was particularly prominent in areas where shallow marine habitats (<60 m) are abundant, such as the Western Atlantic, Western Pacific, and Eastern Indian Ocean [25]. Map projections of the TEP during the LGM show relatively small differences of the exposed landmasses at low sea level (−60m) compared to present day [25], possibly indicating that glaciation sea level drops might not have changed the overall topology and gene flow in the TEP as much as it did in other ocean basins. Overall, although our demographic estimates varied considerable with our choice of mutation rate, our results are generally consistent with previous studies indicating that population expansions of marine fishes in the TEP may have preceded the LGM [29]. Furthermore, this also suggests that the demography history in *H. passer* is likely to be shaped by historical events associated with the closure of the Isthmus of Panama, rather than by the more recent LGM.

## Methods

### Sample collection and DNA extraction

Fin and gill clips were collected from 13 individuals of *Holacanthus passer* in La Paz, Baja California Sur, Mexico. Collections were made with pole spears while SCUBA diving, abiding by IACUC protocols. Tissue samples were immediately placed in 95% ethanol and stored at −20°C. DNA was extracted using a DNeasy Blood and Tissue kit according to manufacturer’s protocol (Qiagen). DNA quality and concentration of the 13 samples were assessed using a Nanodrop 2000c and Qubit 4.0 Fluorometer. The sample with the highest quality was further evaluated on an Agilent 2200 TapeStation DNA ScreenTape to check for high molecular weight. The sample chosen to carry out the genome assembly of *Holacanthus passer* had a final DNA concentration of 205 ng/μl, a 260/280 and 260/230 ratio of 2.02 and 2.26, respectively, and an average fragment length of 38 kb (Figure 5A). This sample came from an adult *H. passer* female with a total length size of 20.4 cm. Before beginning with our library prep, DNA was transferred from AE buffer to EB to remove traces of EDTA, as recommended by Nanopore library prep, using a 3X KAPA Pure Bead clean up (Roche Molecular Systems). DNA was then eluted in 90 μl of EB, reaching a final concentration of 128 ng/μl. This sample was sequenced using ONT and Illumina (HiSeq4000; 150 bp paired-end) sequencing.

**Figure 5.**
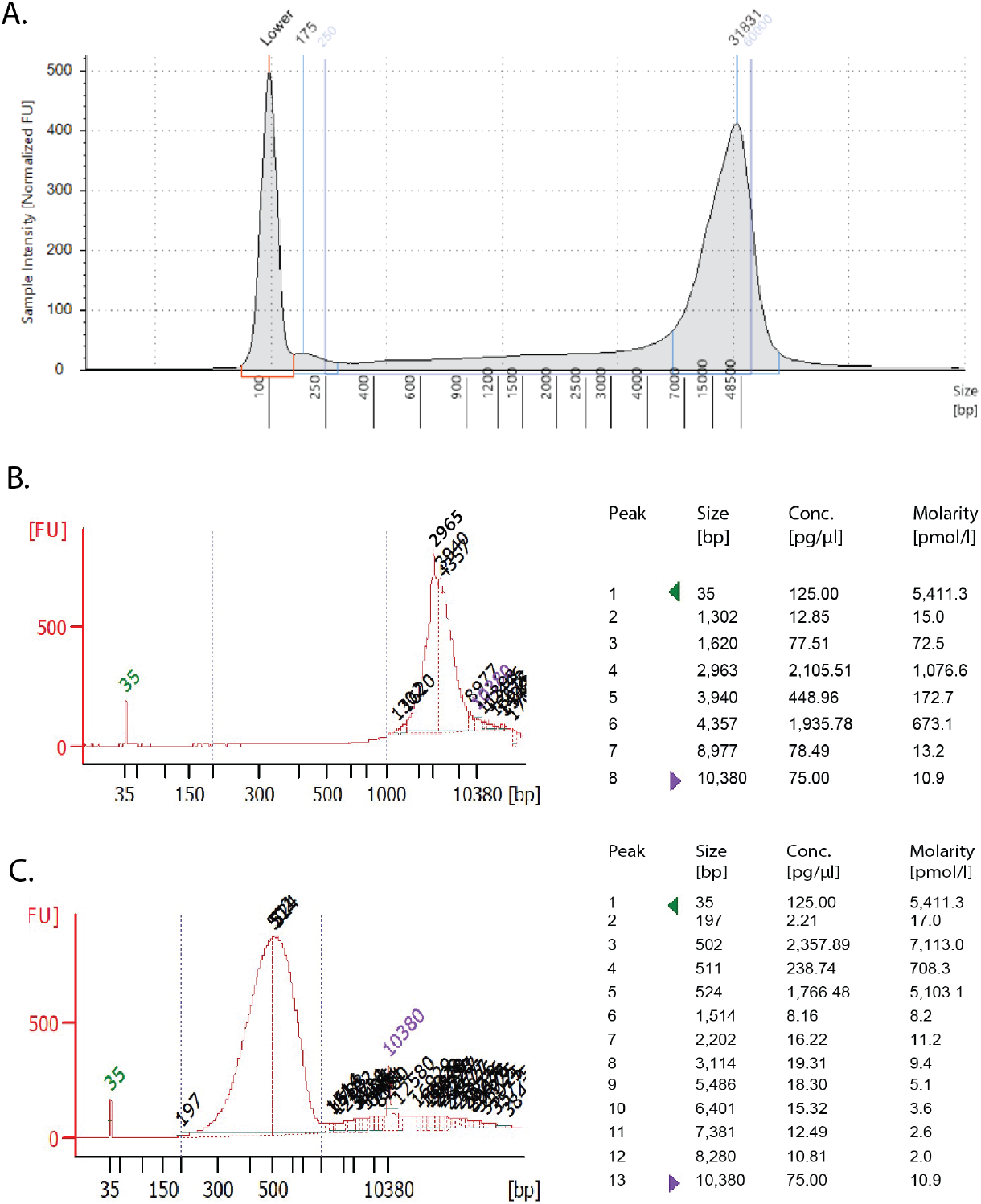
*Holacanthus passer* genomic DNA profile used for Nanopore Sequencing. (A) TapeStation analysis using a Genomic DNA ScreenTape (Agilent Technologies, Inc 2017) of DNA sample used pre-fragmentation. Peak molecular weight was found to be at 31831 bp with a calibrated concentration of 19.6 ng/μl. Between 250 and 60000 bp, a region representing 84% of the sequences, the average size was 18931 bp with a concentration of 23.5 ng/μl. (B;C) Bioanalyzer 2100 profile and statistics using a High Sensitivity DNA Assay (Agilent Technologies, Inc 2009) of genomic DNA post sheared with Covaris g-TUBE following manufacturers protocol for 10 kb fragments (B) and after Kapa Hyperplus library prep followed by a double size-selection cleanup with SPRIselect beads (0.56X and 0.72X)(C).

### Whole-genome library construction and sequencing

Four individual ONT libraries were prepared with 1.5 μg of DNA using the SQK-LSK109 library prep protocol according to manufacturer’s protocol (Oxford Nanopore Technologies, Oxford, UK). DNA was first sheared using the Covaris g-TUBE following the manufacturer’s protocol for 10 kb fragments to improve output yield (Figure 5B). One ONT library was carried out without DNA shearing to target longer fragments, however, N50 only increased about 1 kb while the output yield decreased between half to a third of the yield, thus we opted to continue to shear. Each library was sequenced on a R9.4 flow cell using the MinION DNA sequencer. Maximum run time ranged between 48 to 72 hours. Raw data was basecalled separately using Guppy 3.3 basecaller on a GPU-based high-performance computer cluster server of the University of Massachusetts Boston. A total of 43.8 Gb (N_50_: 6626 bp, longest read: 474 205 bp) were generated on the Oxford Nanopore MinION device. Individual MinION sequencing statistics can be found in the GitHub repository (https://github.com/remygatins/Holacanthus_passer-ONT-Illumina-Genome-Assembly).

The Illumina library was prepared with 250 ng of unsheared DNA using the Kapa Hyperplus Library Preparation Kit with only one third of the volume reactions as described in the manufacturer’s protocol (Kapa Biosystems, Wilmington, MA). The total fragmentation volume was 16.66 μl and was incubated at 37°C for 7:45 min. The incubation parameters were previously optimized to target fragments of ∼500 bp. Post-ligation purification was done using a 0.8X KAPA Pure bead cleanup. Library amplification was carried out with a total PCR reaction volume of 16.6 μl for 8 PCR thermal cycles. Finally, we did a double size-selection post-amplification cleanup with SPRIselect beads using a 0.56X upper and 0.72X lower selection ratio (Beckman Coulter, Inc) (Figure 5C). The final Illumina library was sequenced in a pool of three individuals with a HiSeq4000 (150 bp paired-end) (Novogene Corporation Inc.), which generated a total of 97.3 Gb sequencing data with an average cleaned read of 149 bp.

GenomeScope [34] was used to estimate genome size, repeat content, and heterozygosity across all k-mers (k = 21) previously detected using Jellyfish v2.2.10 [35] to help choose parameters for downstream analysis. Using only raw Illumina data, the genome size of *H. passer* was estimated to have a length of 579 Mb with approximately 95.1% of unique content and a heterozygosity level of 0.43% (Figure 2A). Additionally, k-mers with 110X coverage showed the highest frequency. Considering a genome size of 579 Mb, the output of 43.8 Gb of ONT and 97.3 Gb of Illumina reads represented a total of 75X and 167X coverage respectively, based on the size of our final genome assembly.

### Genome assembly

Long reads obtained from the ONT were concatenated into one large fastq file and trimmed with Porechop v. 0.2.3 (https://github.com/rrwick/Porechop). Nanofilt v. 2.5.0 (https://github.com/wdecoster/nanofilt) was used to create two different filtered data sets to help the contiguity of the final assembly. Our top five longest reads ranged from 176 kb to 474 kb with an average quality score (Q) of 3.9. Thus, the first data set was filtered to keep sequences with a minimum Q score of 3 and sequence length of 1000 bp as it resulted in the most contiguous assembly (Nanofilt parameters -q 3; -l 1000). For the second data set we increased the Q score to 5 and it was explicitly used for downstream assembly polishing (-q 5 and -l 500). The former sequences were assembled using Wtdbg2 v2.5 [36], setting a minimum sequence length of 1000 bp (-L 1000). To improve the draft assembly, two rounds of consensus correction were performed using the -q 5 filtered ONT reads by mapping reads to the draft genome with Minimap2 v. 2.17 and polishing with Racon v. 1.4.7.

Short accurate Illumina reads were used to further polish the ONT genome. Raw sequences were adapter-trimmed with Trimmomatic v. 0.39 [37] and quality checked before and after trimming using FastQC v 0.11.8 (http://www.bioinformatics.babraham.ac.uk/projects/fastqc/). Two rounds of polishing were carried out by mapping the trimmed short reads to the assembly using BWA v 0.7.17 [38], sorted and indexed with Samtools v 1.9 [39], and consensus corrected using Pilon v 1.23 [40].

Finally, given that the DNA used for the genome assembly was extracted from gill tissue, which could be exposed to microorganisms, the final assembly was screened for sequences of bacteria, viruses, and plasmids using Kraken 2.0.9 [41] and Blobtools2 [42]. Any conataminants found and in accordance with both programs were removed from the final assembly. Genome completeness was assessed using Benchmarking Universal Single-Copy Orthologs (BUSCO v3.0.2) [43,44] by comparing the *H. passer* genome to the Actinopterygii (n = 4,584) and Eukaryota (n = 303) ortholog gene datasets. Assembly statistics and BUSCO completeness were assessed after the initial draft assembly, and subsequently, after each polishing iteration (Table 1). The complete flow chart of the full genome assembly pipeline is shown in Figure 6.

**Figure 6.**
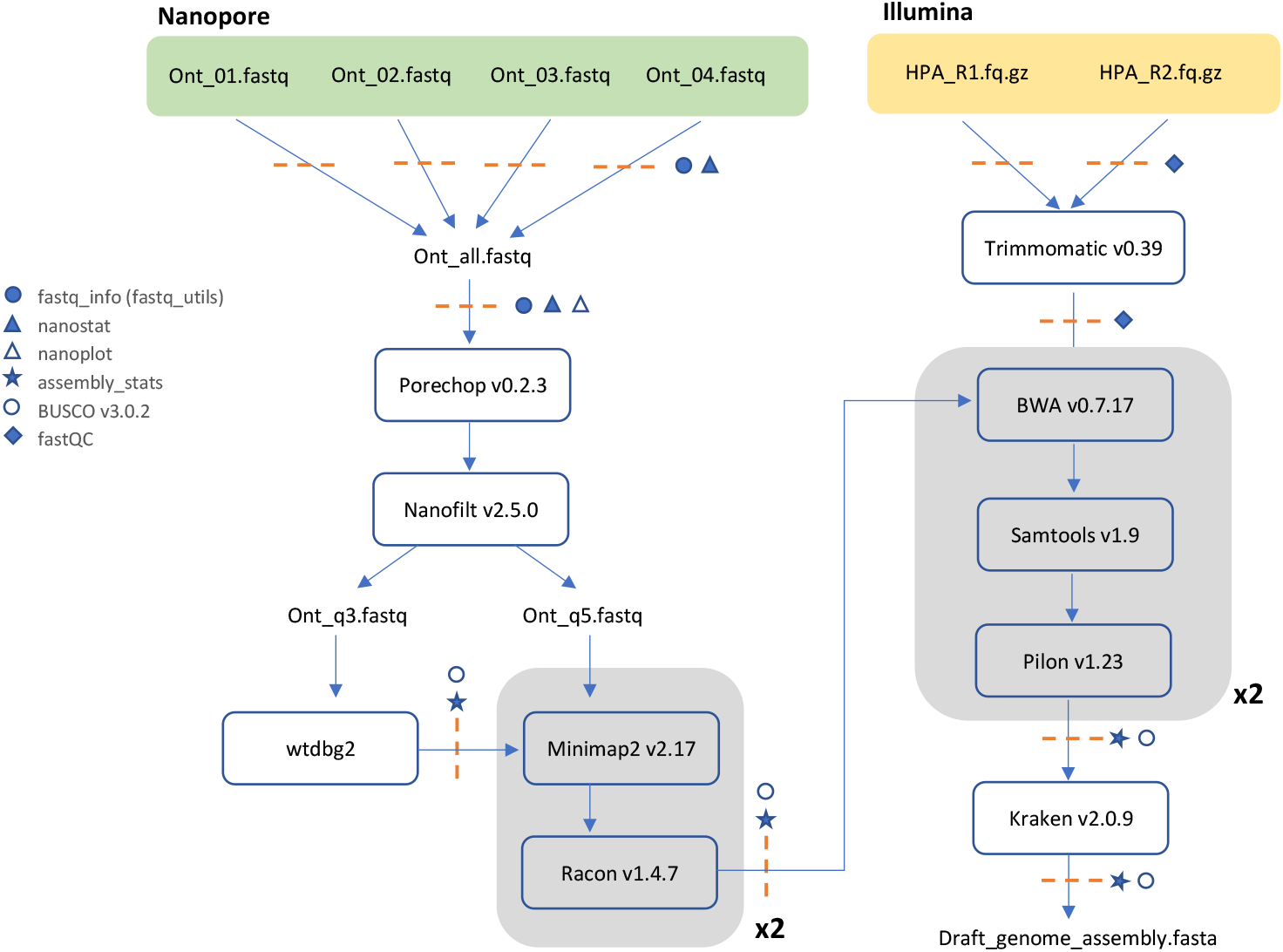
Whole genome assembly pipeline using Oxford Nanopore and Illumina sequencing. Dashed orange lines indicate quality assessment checkpoints carried out during the assembly pipeline.

### Genome annotation

To annotate our genome, we used the homology-based gene prediction pipeline GeMoMa (v1.6.4). GeMoMa uses protein-coding genes models and intron position conservation from reference genomes to predict possible protein-coding genes in a target genome (Keilwagen et al., 2018). Here, we ran the GeMoMa pipeline using annotations from three fish species: *Amphiprion ocellaris, Oreocromis niloticus, Electrophorus electricus* (downloaded from NCBI, see Table 4). These species were selected to represent a variety of genes from close to distant high-quality fish annotations. In our particular case, the pipeline performed four main steps: 1) Extractor or external search, using the search algorithm tbalstn with cds parts as queries from our reference genomes, 2) Gene Model Mapper (GeMoMa), which builds gene models from the extractor results, 3) GeMoMa Annotation Filter (GAF) that filters and combines common gene predictions and 4) AnnotationFinalizer, which predicts UTRs for annotated coding sequences and generate genes and transcripts names (Keilwagen et al., 2018). Additionally, repetitive elements were predicted by running RepeatMasker (open-4.0.6, Smit et al. 2013–2015) with the Teleostei database to identify repetitive elements in the genome and soft-mask the assembly. RepeatMasker.out was converted to GFF with RepeatMasker script ‘rmOutToGFF3.pl’.

**Table 4.** Reference Genomes and annotations used to predict gen models with GeMoMa pipeline.

| Common Name | Scientific Name | RefSeq Assembly | Genome and Annotation Release link | Download date | Annot. Release |
| --- | --- | --- | --- | --- | --- |
| Electric Eel | *Electrophorus electricus* | fEleEle1.pri (GCF_013358815.1) | https://ftp.ncbi.nlm.nih.gov/genomes/all/annotation_releases/8005/101/GCF_013358815.1_fEleEle1.pri/ | 12/1/20 | 101 |
| Clown Anemonefish | *Amphiprion ocellaris* | AmpOce1.0 (GCF_002776465.1) | https://ftp.ncbi.nlm.nih.gov/genomes/all/annotation_releases/80972/101/GCF_002776465.1_AmpOce1.0/ | 12/1/20 | 101 |
| Nile Tilapia | *Oreochromis niloticus* | O_niloticus_UMD_NMBU (GCF_001858045.2) | https://ftp.ncbi.nlm.nih.gov/genomes/all/annotation_releases/8128/104/GCF_001858045.2_O_niloticus_UMD_NMBU/ | 12/1/20 | 104 |

### Demographic history of H. passer

To infer the demographic history of *H. passer* in the TEP, a Pairwise Sequentially Markovian Coalescent (PSMC) model was used to explore temporal changes in effective population size based on genome-wide diploid sequence data [45]. The PSMC analysis is particularly powerful to infer demographic histories beyond 20,000 years, which fits well with the known history of the *Holacanthus* genus [7,8]. The PSMC simulation was run with 30 iterations (-N), a maximum 2N0 coalescent time of 30 (-t), initial theta/rho ratio of 5 (-r), and the pattern parameter (-p) set to “4+30*2+4+6+10” [45,46]. Generation time (g) is defined as the age at which half of the individuals of the population are reproducing. Given that *H. passer* is protogynous, generation time for females is around three years, while for males it is around six years, after they transition from female to male [2,47,48]. Thus, we set the average generation time (-g) for *H. passer* to 5 years. Mutation rate (μ) per site per generation in fishes has been estimated to be between 10^−8^ to 10^−9^ mutations per site [22,49], thus we ran two simulations to represent the range of the expected mutation rates.

## Reuse potential

This study presents the first annotated genome assembly of the King Angelfish, *Holacanthus passer.* It will serve as a resource to improve our understanding of the evolution of *Holacanthus* angelfishes while facilitating novel research into local adaptation, speciation, and introgression in marine fishes. In addition, this genome will help our understanding of the evolutionary history and population dynamics of marine species in the Tropical Eastern Pacific.

## Data Availability

The genome assembly and raw sequencing reads (Illumina and Nanopore) have been deposited into NCBI under BioProject PRJNA713824 and are linked to Biosample SAMN18269499. The GenBank accession number of the genome assembly is JAFREQ000000000.1. Genome annotation and any additional annotation files can be found in Dryad https://doi.org/10.7291/D1X10B. Step-by-step code to reproduce the methods can be found at https://github.com/remygatins/Holacanthus_passer-ONT-Illumina-Genome-Assembly

## List of Abbreviations

bp: base pair
BUSCO: Benchmarking Universal Single-Copy Orthologs
g: Generation time
Gb: gigabase
kb: kilobase
Kya: Thousand years ago
LGM: Last Glacial Maximum
Mya: Million years ago
ONT: Oxford Nanopore
PSMC: Pairwise Sequentially Markovian Coalescent model
TEP: Tropical Eastern Pacific
TWA: Tropical Western Atlantic
μ: Mutation rate

## Declarations

### Ethics approval

The care and handling of all vertebrate animals used in this study was in compliance with the Institutional Animal Care and Use Committee (IACUC/BERNG-1601) of the University of California Santa Cruz and performed in accordance with the Molecular Ecology and Evolution of Fishes Laboratory.

### Consent for publication

Not applicable

### Author Contributions

R.G., C.F.A, C.S., G.B., and L.F.D. designed the project. R.G. and C.S. collected the samples. R.G. and C.F.A carried out the molecular lab work and bioinformatic analyses. All authors contributed to writing and revising the manuscript.

## Acknowledgements

We would like to thank the lab members from Proyecto de Fauna Arrecifal lab at the Universidad de Baja California Sur, La Paz (UABCS) for helping secure fieldwork logistics and sample collections. We would additionally like to thank the UMass Boston High Performance Computing team for their assistance with our computational research needs.

